# Maternal cortisol is associated with neonatal amygdala microstructure and connectivity in a sexually dimorphic manner

**DOI:** 10.1101/2020.06.16.154922

**Authors:** David Q Stoye, Manuel Blesa, Gemma Sullivan, Paola Galdi, Gillian J Lamb, Gill S Black, Alan J Quigley, Michael J Thrippleton, Mark E Bastin, Rebecca M Reynolds, James P Boardman

**Author notes:** Equal contributions (Rebecca M Reynolds, James P Boardman). **Corresponding author**: Professor James P Boardman, MRC Centre for Reproductive Health W1.26 Queen’s Medical Research Institute, 47 Little France Crescent, Edinburgh, EH16 4TJ, UK. E.

## Abstract

The mechanisms linking maternal stress in pregnancy with infant neurodevelopment in a sexually dimorphic manner are poorly understood. We tested the hypothesis that maternal hypothalamic-pituitary-adrenal axis activity, measured by hair cortisol concentration, is associated with microstructure, structural connectivity and volume of the infant amygdala. In 78 human mother-infant dyads, maternal hair was sampled postnatally, and infants underwent magnetic resonance imaging at term-equivalent age. Higher hair cortisol concentration was associated with higher left amygdala fractional anisotropy (β=0.677, p=0.010), lower left amygdala orientation dispersion index (β=-0.597, p=0.034), and higher fractional anisotropy in connections between the right amygdala and putamen (β=0.475, p=0.007) in girls compared to boys. Maternal cortisol during pregnancy is related to newborn amygdala architecture and connectivity in a sexually dimorphic manner. Given the fundamental role of the amygdala in the emergence of emotion regulation, these findings offer new insights into mechanisms linking maternal stress with adverse neuropsychiatric outcomes of children.

**Impact Statement:** Prenatal stress is transmitted to infant development through cortisol, which imparts sex-specific effects on the development and connectivity of the amygdalae.

## Introduction

Prenatal exposure to maternal stress is estimated to affect 10-35% of children worldwide, which is a major concern because early life stress is linked to impaired cognitive development, negative affectivity, autism spectrum disorder (ASD), and psychiatric diagnoses including attention deficit hyperactivity disorder (ADHD), addiction, depression and schizophrenia(1). Neural correlates of prenatally stressed children include alternations in brain structural and functional connectivity, especially in networks involving the amygdala and prefrontal cortex(2).

Adaptation of the maternal hypothalamic-pituitary-adrenal (HPA) axis is a key mechanism by which maternal stress modulates offspring neurodevelopment(3), and there is evidence that this mechanism operates in a sexually dimorphic manner(4). For, example, higher waking maternal salivary cortisol in pregnancy is associated with increased internalizing behaviours in female infants and reduced internalizing behaviours in males(5, 6). Higher maternal salivary cortisol in pregnancy is also associated with stronger amygdala functional connectivity with networks involved in sensory processing and integration in newborn girls, with weaker connectivity to these brain regions in boys(7); and in childhood, with larger amygdalae(8) and reduced segregation of structural networks in girls but not boys(9). The amygdala is further implicated as a neural target of prenatal stress exposure by observations from studies that have characterised maternal stress by symptomatology of depression and / or anxiety, which report alterations in amygdala volume(10), microstructure(11), and functional and structural connectivity among offspring(12)

Candidacy of the amygdala as an important neural target of prenatal stress exposure comes from the following observations in pre-clinical and clinical studies. First, the amygdala develops early in embryonic life(13) and contains a high concentration of glucocorticoid receptors(14); second, increased maternal glucocorticoids modulate amygdala development and anxiety-like behaviours in experimental models(15, 16); third, lesion studies in non-human primates support its critical role in early development of emotion regulation(17); fourth, newborn amygdala functional connectivity is consistently linked with internalizing behaviours in children up to the age of two years(7, 18); fifth, early disruption to cell composition of the amygdala is reported in a model of early life stress(19), and in children with autism(20); and sixth, in pre-clinical models, stress and glucocorticoid exposure induce dendritic arborization, amygdala hypertrophy and induce anxiety-like behaviours(21, 22).

Neonatal magnetic resonance imaging (MRI) serves as an intermediate phenotype for investigating the impact of early life exposures on brain and health because it is distal to the aetiological process, in this case prenatal stress, and is also more proximal to cognitive, behavioural and disease outcomes. Structural and diffusion MRI (dMRI) have been used to characterise brain structural maturation and emerging network connectivity during the perinatal period, and to investigate pathways to atypical development(23, 24). It is a suitable tool to investigate the impact of prenatal stress exposure on the amygdala because age-specific templates enable accurate parcellation of the amygdala and associated structures(25); and diffusion tensor imaging and neurite orientation and dispersion density imaging (NODDI) support inference about tissue microstructure and network connectivity, modelled by fractional anisotropy (FA), mean diffusivity (MD), orientation dispersion index (ODI) and neurite density index (NDI)(26).

Hair cortisol concentration (HCC) measured in 3cm hair samples collected from close to the scalp reflects basal HPA axis activity over the 3 months prior to sampling, and in contrast to single measures from saliva or blood, it is not influenced by short-term activation of the HPA axis in response to acute stressors(27). Studies in pregnant women have shown HCC to be an efficient method of retrospective assessment of long-term cortisol secretion, and thus long-term HPA axis activity(28, 29).

Previous studies have reported sex-specific differences between maternal stress and amygdala functional connectivity and behavioural outcomes among children(5–9), but study designs leave uncertainty about the mechanism linking maternal stress with amygdala development, the potential confounding role of events and environmental exposures during childhood, and the impact of stress on structural connectivity. Resolving these uncertainties is necessary for developing strategies designed to improve socio-emotional development of children born to women who are stressed during pregnancy. Based on studies of the imaging, biochemical and clinical phenotype of prenatal stress exposure, we hypothesised that higher levels of maternal HPA activity in the final months of pregnancy ascertained from maternal HCC would impact amygdala development and structural connectivity of offspring infants in a sexually-dimorphic manner, and that these effects would be apparent around the time of birth.

## Results

### Participant characteristics

The parents of 102 infants consented to take part. Of these, 2 preterm infants died before term equivalent age, 12 did not complete the MRI protocol or images were not amenable processing due to movement artefact; 1 had an incidental structural anomaly detected at MRI; and 9 withdrew before MRI scan. This left data from 78 mother-infant dyads for analysis, the maternal and infant characteristics for whom are shown in Table 1. Maternal hair was sampled at mean 3.5±2.5 days after delivery, and the median HCC concentration was 5.6pg/mg (0.5-107.1). Maternal HCC was not associated with gestational age (GA) at birth (r=0.200, p=0.094). HCC did not differ between mothers of male and female infants (p=0.997). MRI was carried out at term-equivalent age: median 41.9 weeks’ GA (range 38.6-45.9).

**Table 1.** Maternal and neonatal characteristics.

| Maternal characteristics, n = 71 |  |
| --- | --- |
| Age (years) | $33.1 \pm 5.2$ |
| BMI ($\text{kg/m}^2$) | $25.2 \pm 4.2$ |
| Primiparous (%) | 41 (58%) |
| Multiparous (%) | 30 (42%) |
| SIMD 2016 quintile n (%)<sup>†</sup> |  |
| 1 | 4 (6%) |
| 2 | 14 (20%) |
| 3 | 10 (14%) |
| 4 | 14 (20%) |
| 5 | 29 (41%) |
| Tobacco smoked during pregnancy, n (%) | 5 (7%) |
| No tobacco smoked during pregnancy, n (%) | 66 (93%) |
| Gestational diabetes (%) | 1 (1%) |
| Preeclampsia (%) | 4 (6%) |
| Receiving pharmacological treatment for depression | 3 (4%) |
| Infant characteristics, n = 78 |  |
| Birthweight (grams) | 2895 (454-4248) |
| Birth weight z-score<sup>‡</sup> | $0.2 \pm 1.1$ |
| Birth gestation (weeks) | 38.4 (24.0-42.0) |
| Male n (%) | 44 (56%) |
| Female | 34 (44%) |
| European ancestry n (%) | 68 (87%) |
| Other | 10 (13%) |
| Singleton n (%) | 63 (81%) |
| Twin | 15 (19%) |

Normally distributed data is presented as mean ± SD. Non-normally distributed data is presented as median (range). <sup>†</sup>Scottish Index of Multiple Deprivation (SIMD) 2016 quintile. First quintile indicates most deprived, and fifth quintile the least deprived. <sup>‡</sup>Calculated according to INTERGROWTH-21<sup>st</sup> standards

### Amygdala microstructure

In univariate analysis, there were moderately strong correlations between both FA and MD, and GA at birth and age at scan (r=0.41-0.64), and weak correlations with birth weight z-score and Scottish 2016 quintile (r=0.24-0.30). There were no significant correlations between FA and MD in amygdalae and ethnicity or infant sex, or maternal parity, age or BMI. There were moderate to strong correlations between NDI in the amygdalae and GA at birth and age at scan (r=0.43-0.74), and a weak correlation with Scottish Index of Multiple Deprivation (SIMD) 2016 quintile (r=0.23-0.26). Weak to moderate correlations were observed between ODI in amygdalae with GA at birth and age at scan (r=0.28-0.42), (Supplementary Table 1).

In multiple linear regression models, there was a significant interaction effect between maternal HCC and infant sex in left amygdala FA (p=0.010) and ODI (p=0.034), with higher maternal HCC being associated with higher left amygdala FA and lower ODI in girls compared to boys (Table 2). When we stratified by sex, there were associations between maternal HCC and infant amygdala microstructure in boys, but not girls. Table 3 shows that in boys, higher maternal HCC was associated with lower left amygdala FA (β=-0.339), lower right amygdala FA (β=-0.287) and NDI (β=-0.215), and higher right amygdala MD (β=0.264) and ODI (β=0.309), after FDR correction.

**Table 2.**
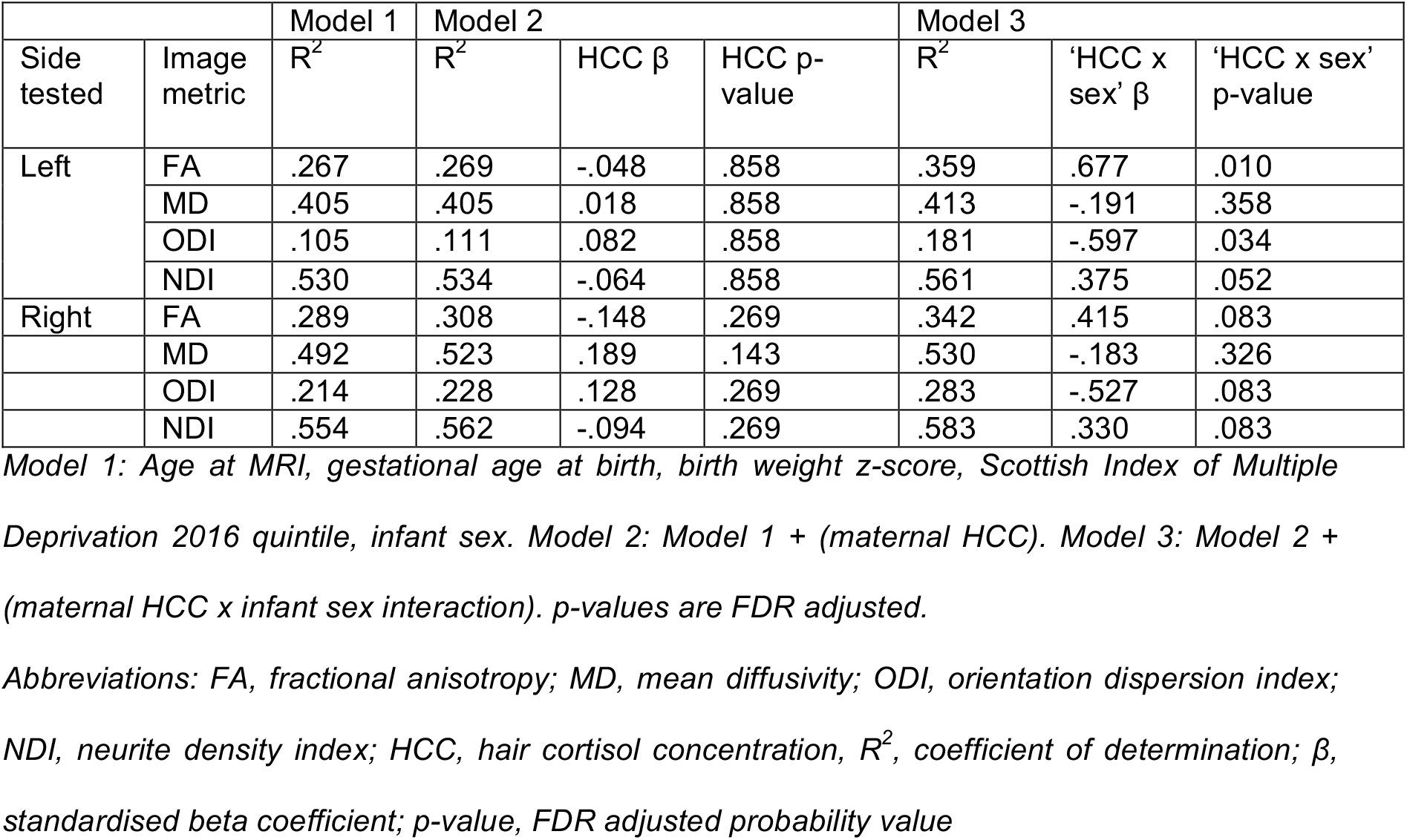
Associations of maternal hair cortisol concentration (HCC) and its interaction with infant sex on amygdala microstructure.

**Table 3.** Associations of maternal hair cortisol concentration (HCC) with amygdala microstructural parameters assessed separately in boys and girls.

|  |  | Boys |  |  |  | Girls |  |  |  |
| --- | --- | --- | --- | --- | --- | --- | --- | --- | --- |
|  |  | Model 1 | Model 2 |  |  | Model 1 | Model 2 |  |  |
| Side tested | Image metric | R<sup>2</sup> | R<sup>2</sup> | HCC $\beta$ | HCC p-value | R<sup>2</sup> | R<sup>2</sup> | HCC $\beta$ | HCC p-value |
| Left | FA | .433 | .537 | -.339 | .023 | .157 | .239 | .340 | .372 |
|  | MD | .489 | .497 | .090 | .462 | .445 | .449 | -.072 | .667 |
|  | ODI | .124 | .215 | .317 | .085 | .132 | .159 | -.194 | .648 |
|  | NDI | .622 | .654 | -.189 | .089 | .522 | .530 | .109 | .648 |
| Right | FA | .368 | .443 | -.287 | .047 | .301 | .306 | .091 | .736 |
|  | MD | .508 | .571 | .264 | .047 | .497 | .506 | .111 | .736 |
|  | ODI | .149 | .236 | .309 | .047 | .362 | .378 | -.149 | .736 |
|  | NDI | .581 | .623 | -.215 | .047 | .571 | .573 | .050 | .736 |

Model 1: Gestational age at MRI, gestational age at birth, birth weight z-score, Scottish Index of Multiple Deprivation 2016 quintile. Model 2: Model 1 + (maternal HCC). p-values are FDR adjusted. Abbreviations: FA, fractional anisotropy; MD, mean diffusivity; ODI, orientation dispersion index; NDI, neurite density index; HCC, hair cortisol concentration; R<sup>2</sup>, coefficient of determination; $\beta$ , standardised beta coefficient; p-value, FDR adjusted probability value

### Structural connectivity of the amygdala

For both hemispheres, the networks with the top 20% number of streamlines were connected to eight structures: thalamus, putamen, insula, superior temporal gyrus, inferior temporal gyrus, middle temporal gyrus, caudate and lateral orbitofrontal cortex, Figure 1. Quantification of streamline counts is given in Supplementary Table 2 and illustrated in Figure 2. Maternal HCC was not associated with streamline counts of the left and right amygdala with these regions.

**Figure 1.**
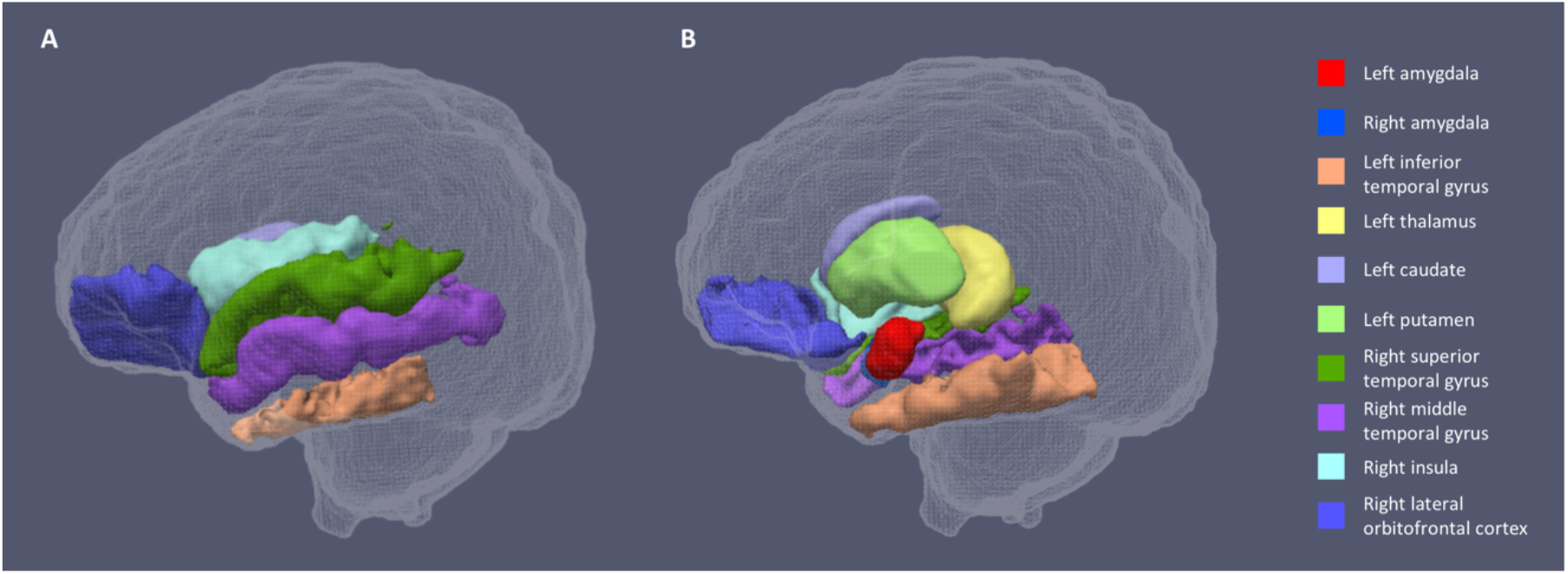
Segmentations of the amygdalae and connected regions defined by the top 20% streamline counts. Figure 1a shows the lateral view of the sagittal plane and 1b the medial view. The same eight regions had the highest streamline counts to the amygdalae bilaterally.

**Figure 2.**
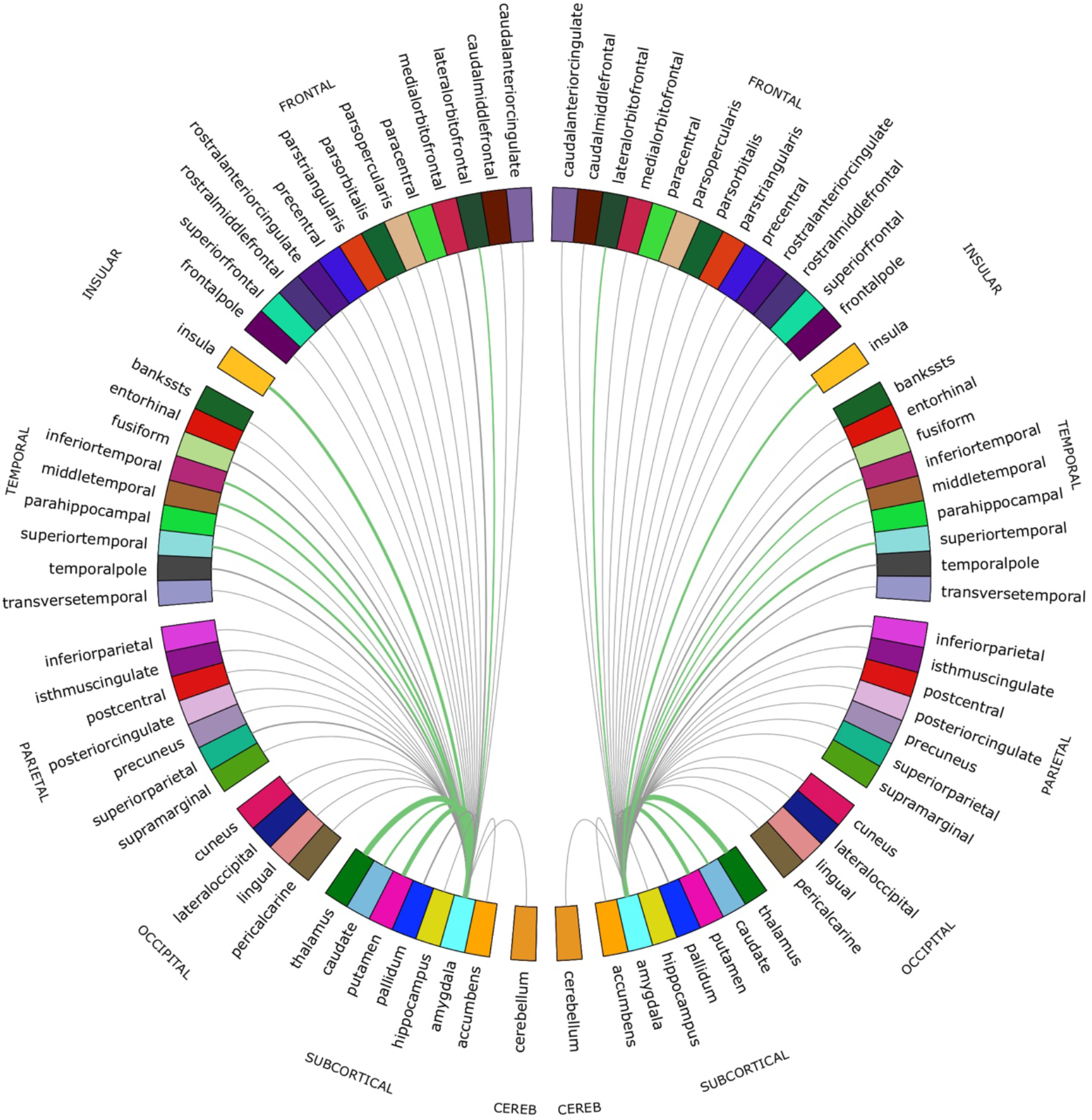
Chord diagram of the streamline counts between the amygdalae and unilateral regions of interest (ROIs). The number of streamlines between ROIs are demonstrated by the corresponding arcs thickness. ROIs connected by the top 20% of streamlines are shown in green.

In fully adjusted analyses, the interaction between maternal HCC and infant sex was significant for mean FA of connections between the right amygdala and putamen. Higher maternal HCC was associated with higher FA for amygdala-putamen connectivity in girls compared with boys (p=0.007). The interaction was also seen for connections to left thalamus, putamen and insula, but the interaction term did not remain after correction for multiple tests (Supplementary Table 3). In sex-stratified analysis, girls had higher FA values in association with high maternal HCC in connections between left amygdala with thalamus, putamen and inferior temporal gyrus, and the right amygdala with putamen and inferior temporal gyrus, but these were not significant after correction for multiple tests (Supplementary Table 4).

### Amygdala volume

Mean volumes of the left and right amygdala were 877±111 mm^3^ and 823±91 mm^3^, respectively. In univariate analysis, there were weak associations (r=0.24-0.3) between amygdala volume and GA at birth and birth weight z-score, but not with age at scan, SIMD 2016 quintile, sex, ethnicity, or maternal BMI, parity or age (Supplementary Table 1). Maternal HCC was not associated with infant right or left amygdala volume in regression models adjusted for potential covariates, and interaction terms between maternal HCC and infant sex were not significant (Supplementary Table 5).

### Sensitivity and sub-group analyses

There were seven twin sets in the whole sample. When we repeated analyses including only singletons and the first born of twin pairs, significant associations between maternal HCC, sex and image feature remained, with little change to the value of regression coefficients (Supplementary Table 6).

In subgroup analysis of preterm and term infants, the direction and magnitude of interaction effects for both groups were similar to those of the whole sample. Specifically, when tested in term and preterm infants, respectively, higher maternal hair cortisol concentration is associated with higher left amygdala fractional anisotropy (β=0.735 and 0.640), lower left amygdala orientation dispersion index (β=−0.710 and −0.614), and higher fractional anisotropy in connections between the right amygdala and putamen (β=0.733 and 0.426) in girls compared to boys (see Supplementary Table 6).

## Discussion

We report a mechanism that could explain the impact of maternal stress on infant brain development. We found that maternal HCC, a stable marker of chronic maternal HPA axis activity in pregnancy, is associated with microstructure and structural connectivity of the newborn amygdala, a region of functional importance for early social development and emotion regulation. Specifically, HCC interacts with infant sex to modify amygdala FA, ODI and NDI, which supports the inference that maternal chronic HPA activity has an impact on dendritic structure, axonal configuration, and the packing density of neurites, in a sexually dimorphic manner (30–33).

The findings are consistent with recent reports from the GUSTO (Growing Up in Singapore Towards Health Outcomes) cohort that describe associations between prenatal depression and alterations in offspring amygdala development(10, 11). That study highlighted the role of maternal mental health on newborn brain development, and focussed attention on the amygdala. Here, we provide mechanistic insights into the relationship between maternal stress and amygdala development with the use of maternal HCC to characterise chronic HPA activity, and the NODDI model for enhanced inference about tissue microstructure. We chose to measure NODDI parameters for assessing microstructure because ODI and NDI in grey matter appear to be functionally tractable. For example, diffusion markers of dendritic density and arborization in grey matter predict differences in intelligence(34), reduced ODI in grey matter is reported in psychosis and in neurodegenerative disease, and reduced grey matter NDI is reported in Parkinson’s disease, Alzheimer’s disease, autism spectrum disorder, and temporal lobe epilepsy (for review see (33)).

Maternal HCC was also related to structural connectivity of the amygdala in a sex-discordant manner. Higher maternal HCC was associated with higher FA in girls than boys in tracts between right amygdala and putamen. These observations were not explained by differences in streamline counts in relation to maternal HCC. Furthermore, in sex-stratified analysis, there were consistent trends for girls born to women with higher HCC to have higher mean FA between the left amygdala and left thalamus, putamen and inferior temporal gyrus, and between right amygdala and right putamen and right inferior temporal gyrus, although these did not survive FDR correction. During the neonatal period, higher FA in white matter tracts is typically taken to imply microstructural maturation, through increased axon diameter, density or myelination. Therefore, increased mean FA demonstrated in connections between the amygdala and putamen, in girls exposed to higher cortisol, could be interpreted as increased maturation of these connections.

The mechanisms underpinning differences in the relationship between maternal cortisol and neurodevelopment of male and female infants are unknown but could occur due to sex differences in the placental metabolism of glucocorticoids(35), regulation of glucocorticoid receptors(36) and secretion and actions of corticotropin-releasing hormone (CRH)(37).

Strengths of this study are the use of biophysical tissue modelling (NODDI) to enable inference about neurite density and organisation in the amygdala; and use of a data driven approach to investigate amygdala structural connectivity. A second strength is use of maternal HCC to operationalise stress because it is a quantitative stable marker of cortisol secretion that represents HPA activity over 3 months; as such HCC is unlikely to reflect transient stresses that can occur in pregnancy, and it overcomes the problems of diurnal variation that occur with plasma and saliva measurements. To our knowledge this is the first study to investigate a physiological measure of chronic maternal HPA activity with quantitative biomarkers of brain development, and to include infants born very preterm. This was important because preterm birth is associated with both exposure to maternal HPA axis dysregulation(38), and an increased risk of inattention and affective disorders(39). The relationships we describe appear to apply across the whole GA range because GA at birth was included in all regression models that were used to investigate association between maternal HCC and image metrics, and in sub-group analyses the magnitude and direction of ‘HCC x sex’ interaction effects were maintained between term and preterm groups. The study has some limitations: first it was not powered to detect both sex and birth gestation interactions, but this should be considered in future study design. Second, follow-up studies that include measures of socio-emotional development are needed to understand functional consequences of these findings. Finally, the newborn amygdalae are relatively small anatomical regions so could be susceptible to partial volume effects influencing microstructural characteristics. To mitigate this risk, we used an age-specific atlas for segmentation, and excluded voxels with a uiso <0.5.

In conclusion, dMRI and HCC were used to investigate mechanisms underlying the transmission of prenatal stressors on infant development. Maternal HCC in pregnancy is associated with newborn amygdala microstructure and structural connectivity, in a sex-dimorphic manner. These findings reveal that the amygdala, a structure of known importance for child development, is susceptible to variations in the prenatal stress environment, and that cortisol imparts sex specific effects on human fetal neurodevelopment.

## Materials and Methods

### Participants

The ‘Stress Response Systems in Mothers and Infants’ cohort recruited mother-infant dyads from the Royal Infirmary, Edinburgh, between March 2018 and August 2019. It prospectively tests associations of perinatal glucocorticoid exposure with brain development, and early life exposures including preterm birth with infant HPA axis regulation. This study recruited mother-infant dyads with birth at ≤32 completed weeks of gestation, and dyads with birth ≥37 weeks’ gestation. Exclusion criteria were congenital fetal abnormality, chromosomal abnormality or regular maternal corticosteroid use. All women gave written informed consent. Ethical approval was granted by South East Scotland 01 Regional Ethics Committee (18/SS/0006).

### Maternal hair cortisol concentrations (HCC)

Maternal hair was sampled within 10 days of delivery. Hair was cut close to the scalp, at the posterior vertex, and stored in aluminium foil at −20°C. The proximal 3cm of hair were analysed by liquid chromatography-tandem mass spectrometry (LC-MS/MS), at Dresden Lab Service GmbH (Dresden, Germany), using an established protocol(40). Adult hair commonly grows at 1cm/month(41) and thus hair segments represented maternal HPA axis activity over the last three months of pregnancy.

### Demographic and clinical information

Participant demographic information was collected through maternal questionnaire and review of medical records. Collected maternal information included: age at delivery (years), parity (primiparous/multiparous), clinical diagnosis of gestational diabetes, pre-eclampsia, pharmacological treatment for depression during pregnancy, antenatal corticosteroid exposure for threatened preterm birth; body mass index (BMI) calculated at antenatal booking; smoking status defined as having smoked any tobacco in pregnancy; SIMD 2016 quintile rank, a score generated by the Scottish government which measures localities’ deprivation according to local income, employment, health, education, geographic access to services, crime and housing. Infant demographics included whether participants were a singleton or twin, ethnicity, GA at birth (weeks), and birth weight z-score calculated according to intergrowth standards(42).

### Magnetic Resonance Imaging

#### Image Acquisition

Infants underwent MRI at term-equivalent age, at the Edinburgh Imaging Facility, RIE. Infants were fed, wrapped and allowed to sleep naturally in the scanner. Flexible earplugs and neonatal earmuffs (MiniMuffs, Natus) were used for acoustic protection. Scans were supervised by a doctor, or nurse trained in neonatal resuscitation.

A Siemens MAGNETOM Prisma 3 T MRI clinical scanner (Siemens Healthcare Erlangen, Germany) and 16-channel phased-array paediatric head and neck coil were used for acquisition(43). In brief, we acquired 3D T1-weighted MPRAGE (T1w) (acquired voxel size = 1mm isotropic) with TI 1100 ms, TE 4.69 ms and TR 1970 ms; 3D T2-weighted SPACE (T2w) (voxel size = 1mm isotropic) with TE 409 ms and TR 3200 ms; and axial dMRI. dMRI was acquired in two separate acquisitions to reduce the time needed to re-acquire any data lost to motion artefact: the first acquisition consisted of 8 baseline volumes (b = 0 s/mm^2^ [b0]) and 64 volumes with b = 750 s/mm^2^, the second consisted of 8 b0, 3 volumes with b = 200 s/mm^2^, 6 volumes with b = 500 s/mm^2^ and 64 volumes with b = 2500 s/mm^2^; an optimal angular coverage for the sampling scheme was applied(44). In addition, an acquisition of 3 b0 volumes with an inverse phase encoding direction was performed. All dMRI images were acquired using single-shot spin-echo echo planar imaging (EPI) with 2-fold simultaneous multislice and 2-fold in-plane parallel imaging acceleration and 2 mm isotropic voxels; except where stated above, all three diffusion acquisitions had the same parameters (TR/TE 3400/78.0 ms).

Conventional images were reported by an experienced paediatric radiologist (A.J.Q.) using a structured system(45). Images with focal parenchymal injury (defined as posthaemorrhagic ventricular dilatation, porencephalic cyst or cystic periventricular leukomalacia) were not included in the final sample.

#### Image Pre-processing

Diffusion MRI processing was performed as follows: for each subject the two dMRI acquisitions were first concatenated and then denoised using a Marchenko-Pastur-PCA-based algorithm(46); the eddy current, head movement and EPI geometric distortions were corrected using outlier replacement and slice-to-volume registration(47–50); bias field inhomogeneity correction was performed by calculating the bias field of the mean b0 volume and applying the correction to all the volumes(51).

The T2w images were processed using the minimal processing pipeline of the developing human connectome project (dHCP) to obtain the bias field corrected T2w, the brain masks and the different tissue probability maps(52). The mean b0 EPI volume of each subject was co-registered to their structural T2w volume using boundary-based registration(53).

### Tissue segmentation and parcellation

The ten manually labelled subjects of the M-CRIB atlas(25) were registered to the bias field corrected T2w using rigid, affine and symmetric normalization (SyN)(54). Next, the registered labels of the ten atlases were merged using joint label fusion(55), resulting in a parcellation containing 84 regions of interest (ROIs).

### Microstructure and volumetric assessments

Volumes were calculated from ROIs derived in the structural images. ROIs were propagated to the diffusion native space using the previously computed transformation.

To calculate the tensor derived metric, only the first shell was used. NODDI metrics were calculated using the recommended values for neonatal grey-matter of the parallel intrinsic diffusivity (1.25 μm^2^·ms^-1^)(56). The obtained metrics are: neurite density index (NDI), isotropic volume fraction (uiso) and orientation dispersion index (ODI). The mean FA, MD, ODI and NDI were calculated for the left and right amygdalae M-CRIB ROIs, after exclusion of voxels with a uiso <0.5. Voxels with a uiso <0.5 were excluded, in order to minimise partial volume effects(57).

### Network construction and analysis

Tractography was performed using constrained spherical deconvolution(CSD) and anatomically-constrained tractography(58, 59) The required 5-tissue type file, was generated by combining the tissue probability maps obtained from the dHCP pipeline with the subcortical structures derived from the parcellation process. Multi-tissue response function was calculated, with a FA threshold of 0.1. The average response functions were calculated. Then, the multi-tissue fiber orientation distribution (FOD) was calculated(60), and global intensity normalization on the FODs images was performed. Finally, the tractogram was created, generating 10 million streamlines, with a minimum length of 20 mm and a maximum of 200 mm and a cut-off of 0.05 (default), using backtrack and a dynamic seeding(61). To be able to quantitatively assess connectivity, spherical-deconvolution informed filtering of tractograms two (SIFT2) was applied to the resulting tractograms(61). The connectivity matrix was constructed using a robust approach, a 2-mm radial search at the end of the streamline was performed to allow the tracts to reach the GM parcellation(62). The final connectivity matrices were multiplied by the μ coefficient obtained during the SIFT2 process.

These connectomes gave a quantification of the SIFT2 weights (referred to as the streamline counts), and the mean FA of connections, between both the left and right amygdala to 41 unilateral regions of interest defined through M-CRIB parcellation. In order to focus analysis on to amygdala’s most structurally connected areas, these 82 ROIs were thresholded according to the number of streamlines connecting them to the left or right amygdala, with the top 20% (N=16) of connections taken forward for further analysis testing relationships with maternal HCC.

### Statistical analysis

Analyses were performed using IBM SPSS Statistics Version 25 Armonk, NY: IBM Corp. Continuous data are summarised as mean ± SD if they had a normal distribution, and median (range) if skewed. Maternal HCC was positively skewed, and log-10 transformed for analysis. The relationship between maternal HCC with infant characteristics was tested using independent t-test and Pearson’s correlation for categorical and continuous variables, respectively. Associations between maternal HCC with i) left and right amygdala microstructure (FA, MD, NDI, ODI), ii) structural connectivity (number of streamlines and mean FA of connections), iii) amygdalae volumes were tested using multiple linear regression. In all models, image feature was the dependent variable and maternal HCC was an independent variable. Covariates included infant sex and clinical or demographic factors that were correlated with either left or right amygdala microstructure or volume using Pearson’s correlation. Associations with the following were tested: GA at birth, age at scan, birth weight z-score, SIMD2016 quintile, infant ethnicity, infant sex, and maternal parity, BMI and age. Antenatal corticosteroid treatment for threatened preterm birth was not included as a covariate because it was given to n=36 (100%) women in the preterm group, was highly correlated with GA at birth (r=0.958, p<0.001), so its inclusion as a covariate would have introduced multicollinearity in regression analysis. For descriptive purposes correlations of infant and maternal factors considered as potential covariates are described as weak if r< 0.3, moderate if r=0.3-0.7, and strong if r>0.7.

Sex differences in the relationship between maternal HCC and newborn imaging features were assessed by adding an interaction term between maternal HCC and infant sex in the whole group regression model. If a significant interaction was present, sex stratified analysis was conducted independently in boys and girls. Benjamini and Hochberg false discovery rate (FDR) correction was used to adjust p-values for multiple testing. FDR corrections were conducted separately for assessments of left amygdala microstructure (n=4), right amygdala microstructure (n=4), left amygdala connectivity (n=8) and right amygdala connectivity (n=8).

One sensitivity analysis was carried out to assess whether associations between maternal HCC and image features might be enhanced by inclusion of twins. We repeated analysis of features with a significant ‘HCC x sex’ interaction in the whole sample, using only singleton pregnancies and the first-born infant of twin pairs. One sub-group analysis of preterm (GA at birth ≤ 32 weeks) and term infants (GA at birth ≥ 37 weeks) was carried out because the relationship between maternal HCC and infant brain development may be gestation specific.

## Materials and Data Availability

All data generated or analysed during this study are included in the manuscript and supporting files.

## Acknowledgments

The work was funded by Theirworld (www.theirworld.org) and was undertaken in the MRC Centre for Reproductive Health, which is funded by MRC Centre Grant (MRC G1002033). RMR acknowledges the support of the British Heart Foundation (RE/18/5/34216). Participants were scanned in the University of Edinburgh Imaging Research MRI Facility at the Royal Infirmary of Edinburgh which was established with funding from The Wellcome Trust, Dunhill Medical Trust, Edinburgh and Lothians Research Foundation, Theirworld, The Muir Maxwell Trust and many other sources; we thank the University’s imaging research staff for providing the infant scanning.

## Competing interests

The authors have no competing interests to declare.

## Notes

### Competing Interest Statement

The authors have declared no competing interest.

