## Supplementary materials for "Maternal cortisol is associated with neonatal amygdala microstructure and connectivity in a sexually dimorphic manner"

*Equal contributions

**Corresponding author:**

**Supplementary materials contains:**

Tables S1 to S6

**Table S1. Univariate analysis of potential covariates with left and right amygdala volume and microstructure**

| | | Age at MRI | GA at birth | Birth weight z-score | SIMD 2016 quintile | Infant ethnicity | Infant sex | Maternal parity | Maternal BMI | Maternal age |
|---|---|---|---|---|---|---|---|---|---|---|
| Left | Volume | -0.077 | 0.237* | -0.136 | -0.030 | 0.176 | .010 | -0.034 | -0.156 | -0.052 |
| | FA | 0.462** | 0.423** | 0.117 | 0.251* | 0.102 | .113 | -0.116 | -0.138 | 0.039 |
| | MD | -0.616** | -0.413** | -0.102 | -0.271* | -0.066 | -.102 | 0.061 | 0.020 | -0.177 |
| | ODI | -0.200 | -0.296** | -0.180 | -0.141 | -0.058 | -.068 | 0.187 | 0.073 | 0.017 |
| | NDI | 0.721** | 0.431** | 0.194 | 0.263* | 0.118 | .134 | -0.072 | 0.008 | 0.135 |
| Right | Volume | -0.201 | 0.104 | -0.298** | -0.189 | -0.029 | -.093 | -0.131 | -0.054 | -0.131 |
| | FA | 0.453** | 0.430** | 0.298** | 0.203 | 0.161 | .163 | -0.125 | 0.021 | 0.034 |
| | MD | -0.643** | -0.543** | -0.237* | -0.239* | -0.145 | -.081 | 0.046 | 0.029 | -0.160 |
| | ODI | -0.275* | -0.418** | -0.263* | -0.094 | -0.196 | -.076 | 0.163 | -0.018 | -0.029 |
| | NDI | 0.738** | 0.449** | 0.210 | 0.228* | 0.118 | .150 | 0.014 | 0.028 | 0.149 |

Pearson Correlation coefficients. p<0.05*, p<0.01**

Abbreviations: FA, fractional anisotropy; MD, mean diffusivity; ODI, orientation dispersion index; NDI, neurite density index; GA, gestational age; SIMD2016, Scottish Index of Multiple Deprivation 2016.

Coding: Antenatal steroids (mother received antenatal steroids for threatened preterm birth=1, mother did not receive antenatal steroids=0), infant ethnicity (European ancestry=0, other=1), infant sex (male=0 female=1), maternal parity (primiparous=0, multiparous=1)

**Table S2. Streamline counts between amygdalae and atlas regions**

| Region of interest | Left amygdala streamline count | Right amygdala streamline count |
|---|---|---|
| Thalamus | .39010 | .37125 |
| Putamen | .30961 | .25621 |
| Insula | .18792 | .15373 |
| Superior temporal gyrus | .15560 | .13090 |
| Inferior temporal gyrus | .14379 | .09229 |
| Middle temporal gyrus | .14303 | .10985 |
| Caudate | .13979 | .13124 |
| Lateral orbitofrontal cortex | .10408 | .11001 |
| Temporal pole | .07476 | .06859 |
| Fusiform gyrus | .07261 | .06533 |
| Pallidum | .06616 | .05741 |
| Medial orbitofrontal cortex | .06413 | .05036 |
| Superior parietal cortex | .05621 | .04571 |
| Precentral gyrus | .05370 | .04776 |
| Supramarginal gyrus | .05308 | .04392 |
| Inferior parietal cortex | .05262 | .06055 |
| Postcentral gyrus | .05062 | .04391 |
| Superior frontal gyrus | .04645 | .04191 |
| Lateral occipital cortex | .04189 | .03917 |
| Accumbens | .03646 | .03166 |
| Hippocampus | .03570 | .03794 |
| Precuneus cortex | .02633 | .02131 |
| Middle frontal gyrus- rostral division | .02612 | .02138 |
| Lingual gyrus | .02318 | .02391 |
| Cerebellar hemisphere | .02159 | .02634 |
| Transverse temporal cortex | .01939 | .01408 |
| Paracentral lobule | .01606 | .01039 |
| Parahippocampal gyrus | .01353 | .01201 |
| Cingulate cortex-posterior division | .01319 | .01225 |
| Inferior frontal gyrus- pars opercularis | .01276 | .00991 |
| Middle frontal gyrus - caudal division | .01179 | .01236 |
| Cingulate cortex- rostral anterior division | .01131 | .01135 |
| Isthmus division of cingulate cortex | .01052 | .01091 |
| Entorhinal cortex | .00989 | .01104 |
| Pericalcarine cortex | .00985 | .01062 |
| Inferior frontal gyrus- pars orbitalis | .00770 | .00570 |
| Inferior frontal gyrus- pars triangularis | .00729 | .00612 |
| Cuneus cortex | .00622 | .00906 |
| Banks of the superior temporal sulcus | .00531 | .00662 |
| Cingulate cortex- caudal anterior division | .00396 | .00391 |
| Frontal pole | .00131 | .00204 |

Highlighted box shows structures connected by the top 20% streamline counts from the amygdalae.

**Table S3. Associations of maternal hair cortisol concentration (HCC) and its interaction with offspring sex on mean fractional anisotropy (FA) of amygdala networks with high streamline count**

| | | Model 1 | Model 2 | | | Model 3 | | |
|---|---|---|---|---|---|---|---|---|
| Side tested | Region of interest | $R^2$ | $R^2$ | HCC β | HCC p-value | $R^2$ | 'HCC x sex' β | 'HCC x sex' p-value |
| Left | Thalamus | .477 | .477 | -.005 | .971 | .522 | .479 | .097 |
| | Putamen | .475 | .475 | .007 | .971 | .506 | .398 | .104 |
| | Insula | .465 | .465 | -.003 | .971 | .501 | .426 | .104 |
| | Superior temporal gyrus | .531 | .532 | .028 | .971 | .544 | .242 | .299 |
| | Inferior temporal gyrus | .635 | .638 | .052 | .971 | .644 | .183 | .343 |
| | Middle temporal gyrus | .561 | .567 | .084 | .971 | .569 | .103 | .560 |
| | Caudate | .276 | .277 | -.027 | .971 | .285 | .196 | .448 |
| | Lateral orbitofrontal cortex | .636 | .636 | -.026 | .971 | .649 | .255 | .228 |
| Right | Thalamus | .438 | .439 | .034 | .829 | .453 | .267 | .294 |
| | Putamen | .506 | .510 | .068 | .829 | .583 | .606 | .007 |
| | Insula | .507 | .507 | .009 | .924 | .525 | .301 | .220 |
| | Superior temporal gyrus | .585 | .586 | .036 | .829 | .587 | .073 | .674 |
| | Inferior temporal gyrus | .558 | .566 | .096 | .829 | .575 | .211 | .311 |
| | Middle temporal gyrus | .573 | .574 | .032 | .829 | .580 | .174 | .370 |
| | Caudate | .316 | .322 | .087 | .829 | .357 | .417 | .220 |
| | Lateral orbitofrontal cortex | .652 | .654 | -.039 | .829 | .667 | .264 | .220 |

Model 1: Age at MRI, gestational age at birth, birth weight z score, Scottish Index of Multiple Deprivation 2016 quintile, infant sex

Model 2: Model 1 + (maternal HCC)

Model 3: Model 2 + (maternal HCC X infant sex interaction)

Abbreviations: HCC, hair cortisol concentration; $R^2$, coefficient of determination; β, standardised beta coefficient; p-values are FDR adjusted probability values.

**Table S4. Association between maternal hair cortisol concentration (HCC) and fractional anisotropy (FA) weighted connections of the amygdalae in boys and girls**

| | | Boys | | | | Girls | | | |
|---|---|---|---|---|---|---|---|---|---|
| | | Model 1 | Model 2 | | | Model 1 | Model 2 | | |
| Side tested | Region of interest | $R^2$ | $R^2$ | HCC β | HCC p-value | $R^2$ | $R^2$ | HCC β | HCC p-value |
| Left | Thalamus | .579 | .614 | -.197 | .388 | .330 | .438 | .389 | .106 |
| | Putamen | .656 | .677 | -.150 | .388 | .268 | .388 | .410 | .106 |
| | Insula | .532 | .558 | -.168 | .388 | .426 | .461 | .221 | .253 |
| | Superior temporal gyrus | .600 | .607 | -.083 | .709 | .413 | .472 | .288 | .165 |
| | Inferior temporal gyrus | .639 | .641 | -.037 | .825 | .659 | .707 | .262 | .106 |
| | Middle temporal gyrus | .608 | .609 | .016 | .878 | .495 | 541 | .256 | .165 |
| | Caudate | .399 | .402 | -.054 | .825 | .256 | .277 | .172 | .428 |
| | Lateral orbitofrontal cortex | .683 | .691 | -.095 | .643 | .607 | .615 | .107 | .448 |
| Right | Thalamus | .583 | .590 | -.087 | .858 | .279 | .347 | .310 | .168 |
| | Putamen | .526 | .562 | -.200 | .484 | .522 | .613 | .359 | .124 |
| | Insula | .631 | .642 | -.111 | .751 | .390 | .426 | .226 | .259 |
| | Superior temporal gyrus | .620 | .620 | -.004 | .968 | .514 | .533 | .166 | .330 |
| | Inferior temporal gyrus | .533 | .533 | -.005 | .968 | .554 | .623 | .310 | .130 |
| | Middle temporal gyrus | .594 | .596 | -.051 | .858 | .507 | .552 | .252 | .168 |
| | Caudate | .373 | .378 | -.075 | .858 | .350 | .431 | .337 | .149 |
| | Lateral orbitofrontal cortex | .682 | .702 | -.148 | .484 | .603 | .613 | .121 | .394 |

Model 1: Age at MRI, gestational age at birth, birth weight z score, Scottish Index of Multiple deprivation 2016 quintile

Model 2: Model 1 + (maternal HCC).

Abbreviations: HCC, hair cortisol concentration; $R^2$, coefficient of determination; β, standardised beta coefficient; p-values are FDR adjusted probability values.; p-values are FDR adjusted probability values

**Table S5. Associations of maternal hair cortisol concentration (HCC) and its interaction with infant sex on amygdalae volume**

| | Model 1 | Model 2 | | | Model 3 | | |
|---|---|---|---|---|---|---|---|
| Side tested | $R^2$ | $R^2$ | HCC β | HCC p-value | $R^2$ | 'HCC x sex' β | 'HCC x sex' p-value |
| Left | .140 | .168 | -.182 | .124 | .180 | .247 | .315 |
| Right | .220 | .223 | -.056 | .624 | .224 | .078 | .743 |

Model 1: Age at MRI, gestational age at birth, birth weight z score, Scottish Index of Multiple deprivations 2016 quintile, infant sex

Model 2 variables: Model 1 + (maternal HCC)

Model 3: Model 2 + (maternal HCC x infant sex interaction)

Abbreviations: HCC, hair cortisol concentration; $R^2$, coefficient of determination; β, standardised beta coefficient; p-values are unadjusted probability values

**Table S6. Sensitivity and subgroup analyses assessing hair cortisol concentration (HCC) and infant sex interactions**

| Imaging metric assessed | Subgroup analysed | $R^2$ | $R^2$ change attributable to HCC x sex interaction | 'HCC x sex' β | 'HCC x sex' p-value |
|---|---|---|---|---|---|
| Whole group analysis, reported in results, n=78 | | | | | |
| Left amygdala FA | | .359 | .090 | .677 | .002 |
| Left amygdala ODI | | .181 | .070 | -.597 | .017 |
| Mean FA of connections between the right amygdala and putamen | | .583 | .072 | .606 | .001 |
| Analysis including only singletons and first-born twins, n=71 | | | | | |
| Left amygdala FA | | .315 | 0.090 | 0.694 | 0.005 |
| Left amygdala ODI | | .174 | 0.094 | -0.708 | 0.010 |
| Mean FA of connections between the right amygdala and putamen | | .594 | 0.081 | 0.659 | 0.001 |
| Sub-group analysis of term and preterm infants (term n=42, preterm n=36) | | | | | |
| Left amygdala FA | Term | .411 | 0.068 | 0.735 | 0.057 |
| | Preterm | .373 | 0.114 | 0.640 | 0.032 |
| Left amygdala ODI | Term | .182 | 0.063 | -0.710 | 0.115 |
| | Preterm | .268 | 0.105 | -0.614 | 0.054 |
| Mean FA of connections between the right amygdala and putamen | Term | .667 | 0.067 | 0.733 | 0.013 |
| | Preterm | .575 | 0.051 | 0.426 | 0.079 |

Model: Age at MRI, gestational age at birth, birth weight z score, Scottish Index of Multiple deprivation 2016 quintile, infant sex, maternal HCC, maternal HCC x infant sex interaction

Abbreviations: HCC, hair cortisol concentration; $R^2$, coefficient of determination; β, standardised beta coefficient; p-values are unadjusted probability values
